# Dynamic urinary proteomic analysis in a Walker 256 intracerebral tumor model

**DOI:** 10.1101/481697

**Authors:** Linpei Zhang, Yuqiu Li, Wenshu Meng, Yanying Ni, Youhe Gao

**Affiliations:** Department of Biochemistry and Molecular Biology, Beijing Normal University, Gene Engineering Drug and Biotechnology Beijing Key Laboratory, Beijing, China; Department of Pathology, Aviation General Hospital of China Medical University, Beijing, China

**Keywords:** Urine, Proteomics, Brain tumors, Biomarker, Animal model

## Abstract

Patients with primary and metastatic brain cancer have an extremely poor prognosis, mostly due to the late diagnosis of disease. Urine, which lacks homeostatic mechanisms, is an ideal biomarker source that accumulates early and highly sensitive changes to provides information about the early stage of disease. A rat model mimicking the local tumor growth process in the brain was established with intracerebral Walker 256 (W256) cell injection. Urine samples were collected on days 3, 5 and 8 after injection and then analyzed by LC-MS/MS. In the intracerebral W256 model, no obvious clinical manifestations changes or abnormal MRI signals were found on days 3 and 5; at these time points, nine proteins were changed significantly in the urine of all 8 tumor rats. On day 8, when tumors were detected by MRI, twenty-five differential proteins were identified, including 10 proteins that have been reported to be closely related to tumor metastasis or brain tumors. The differential urinary proteomes were compared with those from the subcutaneous W256 model and the intracerebral C6 model. Few differential proteins overlapped. Specific differential protein patterns were observed among the three models, indicating that the urinary proteome can reflect the difference when tumor cells with different growth characteristics are inoculated into the brain and when identical tumor cells are inoculated into different areas, specifically, the subcutis and the brain.

## Introduction

Brain tumors can be divided into two main types: primary brain tumors that start within the brain, such as astrocytoma, and brain metastases, which spread from a distant primary tumor in another organ and are the most common intracranial tumor in adults[1, 2]. Brain metastases occur for approximately 20-40% of malignant tumors, and the main metastases arise from lung cancer, breast cancer and melanoma[3, 4]. The prognosis of malignant brain tumors is extremely poor, and the overall survival of patients is significantly shortened, which seriously affects the quality and length of cancer patients’ life[5]. Therefore, there is an urgent need for biomarkers that can detect brain tumors, thereby enabling reasonable and effective treatment measures for patients.

An important biomarker source, urine, is not regulated by homeostatic mechanisms, can reflect changes in the whole body, and can sensitively reflect changes caused by lesions of various organs, even the brain, which has the blood-brain barrier. Some studies showed that urine has the potential to contain biomarkers for brain diseases[6]. Additionally, urine proteomics is an exciting field that provides a tool for tumor marker discovery, and new developments have occurred in recent years. When animal models are used to evaluate urine markers, the start point and the entire development process of the tumor can be controlled; thus, early-stage urine samples can easily be collected. Moreover, the influence of genetic background, living environment, and drugs on clinical urine samples can be circumvented.

We recently demonstrated that urinary proteins could enable the early detection of cancer in a Walker 256 (W256) tumor-bearing model using a urinary proteomics approach[7]. In a glioma rat model injected with C6 cells, changes in urinary proteins were found earlier than magnetic resonance imaging (MRI) changes[8]. Therefore, we asked whether inoculating the same tumor cells into different tissues would cause different urinary changes and whether inoculated different tumor cells in the same organ would cause different urinary proteome changes.

In this study, a rat model was established by intracerebral injection of W256 breast carcinoma cells. Magnetic resonance imaging, the most widely used technique for diagnosing brain diseases in the clinic, was used to monitor the tumor growth[9]. The urinary proteome in rats was analyzed after intracerebral W256 injection with a label-free proteomics method. Then, the differential proteomes of intracerebral W256 model were compared with those from the subcutaneous W256 tumor-bearing model and the intracerebral C6 glioblastoma multiforme model to investigate the ability of the urine proteome to distinguish different tumor lesions.

## Materials and methods

### Animal model

Thirty male Wistar rats (180-200 g) were purchased from Beijing Vital River Laboratory Animal Technology Co., Ltd. The experiment in this study was approved by the Institutional Animal Care Use & Welfare Committee of the Institute of Basic Medical Sciences, Peking Union Medical College (Animal Welfare Assurance Number: ACUC-A02-2014-008). All animals were housed in a standard environment (12 h light/12 h dark cycle, 22 ± 1°C room temperature and 65-70% humidity). The W256 breast carcinoma cell line was obtained from Cell Culture Center of Chinese Academy of Medical Sciences (Beijing, China).

Experimental rats (n=19) were anesthetized with an intraperitoneal injection of a 2% sodium pentobarbital solution at 20 mg/kg and placed in a stereotaxic apparatus. A midline incision was made in the scalp, and a burr hole was drilled above the injection site (bregma: +1 mm; right 3 mm; depth 5 mm). Five microliters of sterile normal saline containing 2000 W256 cells was injected using a 100 µL microsyringe. After injection, the scalp wound was closed with bone wax and sutured. The control rats (n=11) were injected with 5 µL of normal saline. All rats were injected with penicillin sodium postoperatively to prevent postoperative infection.

### Magnetic resonance imaging

On days 2, 5, 7, 9, and 12 after the tumor cell injection, four randomly selected rats in the experimental group were subjected to small animal MRI scans using PharmaScan 70/16 US (Bruker, Switzerland). Rats were anesthetized with 2.5% isoflurane gas. The scan sequence was T2_TurboRARE. The scanning parameters were set as follows: TR/TE=3700/33 ms; slice thickness: 0.5 mm; field of view: 35 × 35 mm; matrix: 256 × 256; number of averages: 3; and acquisition time: 5 min 30 s.

### Histopathology analysis

Rats were perfusion-fixed under anesthesia with normal saline followed by 4% paraformaldehyde via the left ventricle. The brain tissues were removed and postfixed in 10% formalin. Then, the tissues were embedded in paraffin and sectioned, and the pathological changes in the brain tissue after tumor cell injection were evaluated with hematoxylin and eosin (H&E) staining.

### Urine collection and sample preparation

Urine samples were collected on days 3, 5, 8 and 10 after tumor cell injection and then stored at - 80°C. During collection, rats were individually placed in metabolic cages overnight naturally for 10 h to collect urine samples, and no water or food was provided to avoid urine contamination.

Urine samples were centrifuged at 12,000 g for 30 min at 4°C to remove impurities and large cell debris. The supernatants were precipitated with three volumes of prechilled ethanol at -20°C for 2 h. After centrifugation, the precipitates were dissolved in lysis buffer (8 mol/L urea, 2 mol/L thiourea, 50 mmol/L Tris, and 25 mmol/L dithiothreitol [DTT]) and then centrifuged at 12,000 g for 30 min at 4°C. The protein in the resulting supernatant was quantified by the Bradford assay.

### Tryptic digestion

Urinary proteins were digested using the filter-aided sample preparation method[10]. Each 100 µg of protein was loaded onto a 10 kDa filter device (Pall, Port Washington, NY, USA). After sequential washing with UA buffer (8 mol/L urea, 0.1 mol/L Tris-HCl, pH 8.5) and 25 mmol/L NH_4_HCO_3_, the proteins were reduced with 20 mmol/L DTT (Sigma) at 37°C for 1 h and then alkylated with 50 mmol/L iodoacetamide (IAA, Sigma) in the dark for 30 min. Then, the samples were digested with trypsin (1:50 enzyme to protein ratio) at 37°C overnight. The resulting peptides were desalted using Oasis HLB cartridges (Waters, Milford, MA) and then dried by SpeedVac (Thermo Fisher Scientific, USA).

### LC-MS/MS analysis

The digested peptides were acidified with 0.1% formic acid, and 1 µg of peptides was loaded onto a trap column (Acclaim PepMap^®^100, 75 µm × 2 cm, nanoViper C18) and separated by an analytic column (Acclaim PepMap^™^ RSLC 100, 75 µm × 25 cm, 2 µm, nanoViper C18) using an EASY-nLC 1200 HPLC system (Thermo Fisher Scientific, USA). The elution gradient was 5-28% buffer B (0.1% formic acid in acetonitrile, flow rate = 0.3 µL/min) over 90 min. Peptides were analyzed using a Thermo Orbitrap Fusion Lumos Tribrid mass spectrometer (Thermo Fisher Scientific, USA)[11]. The MS data were acquired using the data-dependent acquisition mode. Survey MS scans were acquired by the Orbitrap in the 350-1550 m/z range with a resolution of 120,000. For the MS/MS scan with the resolution set to 30,000, a higher energy collision-induced dissociation (HCD) collision energy of 30 was chosen. Dynamic exclusion was employed with a 30 s window. Two technical replicate analyses were performed for each sample.

Four urine samples in control rats, twenty-four urine samples at three time points in four tumor rats (days 3, 5 and 8) and in four tumor-rejecting rats (days 5, 8 and 10) were chosen for LC-MS/MS analysis. Also, the same method was used for biomarker validation samples except for the different analytic columns (Acclaim PepMap^™^RSLC 100, 50 µm × 15 cm, 2 µm, nanoViper C18) and elution times (60 min).

### Data analysis

Raw data files from the intracerebral W256 model were searched with Mascot Daemon software (version 2.5.1, Matrix Science, UK) against the SwissProt_2017_02 database (taxonomy: Rattus; containing 7,992 sequences) with the following parameters: trypsin digestion was selected, 2 sites of leaky cutting were allowed, and carbamidomethylation of cysteines was set to fixed modification. A peptide mass tolerance of 10 ppm and a fragment mass tolerance of 0.05 Da were applied. Proteins were then filtered using the decoy database method in Scaffold (version 4.7.5, Proteome Software Inc., USA). The proteins were identified with a protein false discovery rate (FDR) < 1%, peptide threshold > 95% and included at least two unique peptides.

The changed urinary proteins were screened with the following criteria: fold change in the increased group ≥1.5, fold change in the decreased group ≤0.67, and P < 0.05 by an independent sample t-test. The protein spectral counts of every rat in the higher group were greater than those in the lower group, and the average fold change of the spectral count in the higher group was ≥2.

In a study with a model other than the intracerebral W256 model, *Wu et al.* analyzed the urine proteome in a tumor-bearing model by subcutaneous W256 injection[7]. Ni *et al.* established a glioblastoma multiforme model by intracerebral injection of C6 cells, and raw data collected by mass spectrometry were reanalyzed using Scaffold Q+ for better and consistent comparisons (detailed information and analysis results are listed in Table S1).

## Results and discussion

### Intracerebral Walker 256 brain metastasis model

The animals displayed no overt clinical signs or weight loss after injection. On day 7, there was a significant difference in body weight between the experimental group and the control group (Figure 1A). On day 9, the tumor inoculation produced consistent tumors in animals, as indicated by MRI analysis, except for four rats that showed no clinical signs or abnormal MRI signal on day 30 after intracerebral injection of W256 cells.

**Figure 1.**
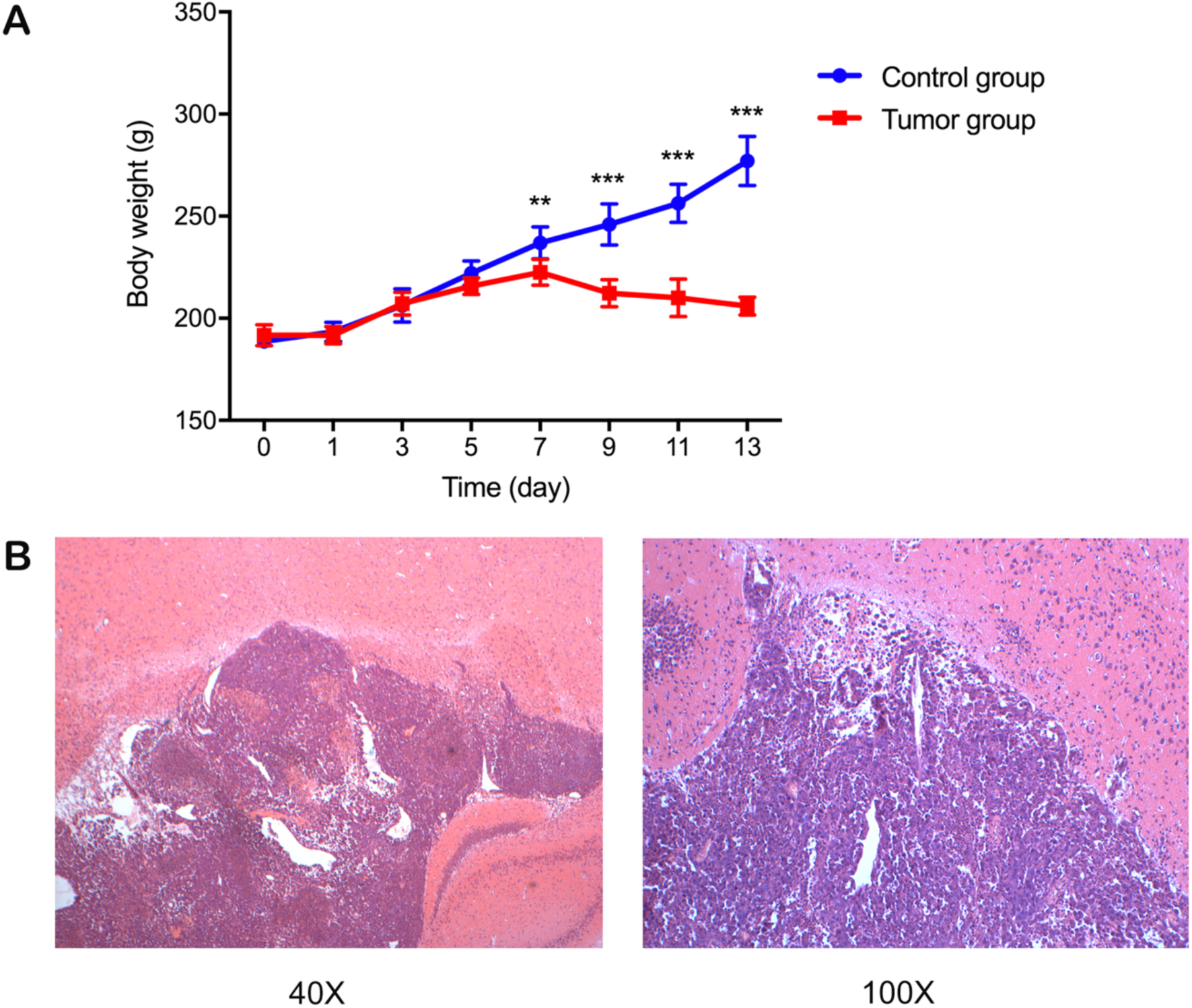
Body weight and pathological changes of the rats injected with W256 cells. A) On day 7, the rats injected with W256 cells exhibited a significantly different body weight than that of the control group and began to show weight loss. P<0.01 (**), P<0.001 (***); B) Pathological changes in brain tissue after W256 cell injection.

The brain tissues of the tumor rats were analyzed by H&E staining for pathological examination. Many tumor cells were observed in the brain tissue, and there was a clear border between the tumor cells and the surrounding brain parenchyma (Figure 1B).

### MRI analysis

On day 5, there was no significant difference in tumor tissue formation between the right side of the brain, which was injected with tumor cells, and the left side of the brain. On day 9, a smaller tumor lesion was detected on T2-weighted images in the right side of the brain, suggesting the presence of a tumor. As the tumor progressed, on day 12, significant tumor-occupying lesions and peritumoral edema were detected on T2-weighted images of the right side of the brain (Figure 2).

**Figure 2.**
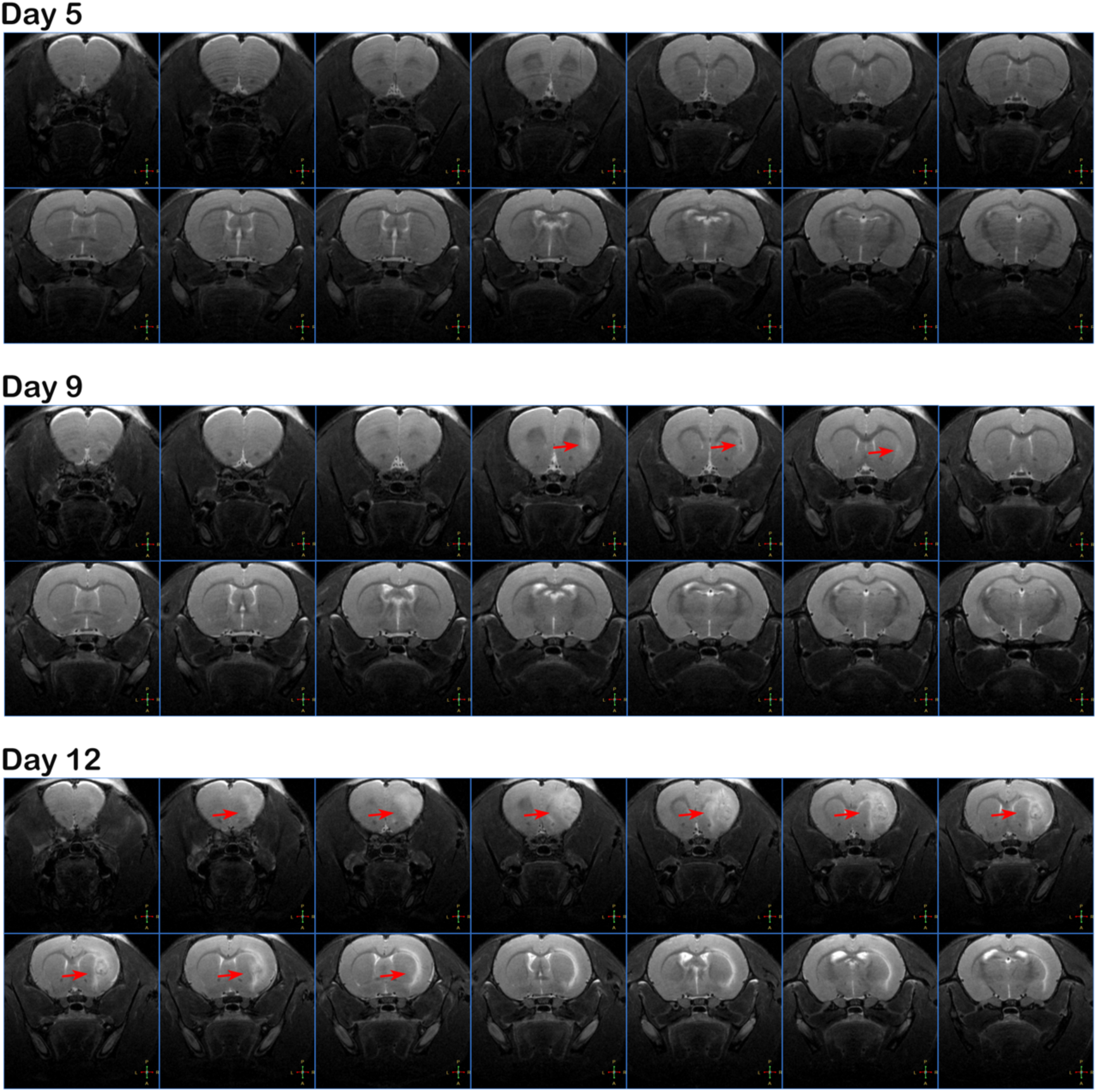
MRI changes in a rat injected with W256 cells.

### Dynamic changes in urinary proteins of intracerebral W256 model rats

The urinary proteomes from sixteen urine samples were profiled by the LC-MS/MS method to investigate the urinary protein changes with tumor progression in the brain. A total of 675 proteins were identified. Among these, 102 proteins were significantly changed in all four rats and included 21, 33, and 75 proteins on days 3, 5, and 8, respectively (fold change ≥1.5 or ≤0.67, P < 0.05).

Until the 5th day after intracerebral injection of W256 cells, the clinical manifestations and MRI images did not significantly differ between the experimental rats and the control rats, but 43 changed proteins were identified in the urine on day 3 and day 5 (Table 1). Gene Ontology (GO) analysis was performed using DAVID 6.8 (https://david.ncifcrf.gov/). Forty out of 43 proteins were annotated and classified into three functional categories: biological process, cellular component and molecular function. Figure 3A shows the 10 most significantly enriched biological processes. Positive regulation of cell migration, the most significant biological process enriched 6 proteins (CSF1, ITGB1, LRRC15, MUC18, PDGFRA and TGFR-1), is a key process for cancer cell dissemination and metastasis. Some proteins, such as CD14, B2MG, NKG2D, TNFR2 and C4, are enriched in the innate immune response, inflammatory response or cellular response to lipopolysaccharide that was associated with body-injected tumor cells. Overexpression of proteins related to negative regulation of extrinsic apoptotic signaling pathway and negative regulation of neuron death may be due to the invasion and proliferation of tumor cells in the brain. In addition, extracellular exosome, extracellular space, cell surface and membrane were the most represented categories in cellular components. The most common molecular functions were receptor binding, protein homodimerization activity, growth factor binding and receptor activity.

**Table 1.** Early-stage changed urinary proteins in the intracerebral W256 model.

| Accession | Protein name | Human ortholog | Day 3 |  | Day 5 |  | Day 8 |  |
| --- | --- | --- | --- | --- | --- | --- | --- | --- |
|  |  |  | FC | P-value | FC | P-value | FC | P-value |
| Q80WY6 | Tumor necrosis factor receptor superfamily member 1B (TNFR2) | P20333 | 6.67 | 5.13E-04 | 8.67 | 1.71E-03 | 10.67 | 1.46E-04 |
| O70215 | NKG2-D type II integral membrane protein (NKG2D) | P26718 | 5.67 | 3.76E-03 | 9.33 | 1.98E-04 | 6.83 | 2.29E-03 |
| P15978 | Class I histocompatibility antigen (RT1AW2) | P01891 | 2.35 | 3.76E-03 | 4.50 | 1.24E-03 | 4.50 | 1.50E-03 |
| O70513 | Galectin-3-binding protein (LG3BP) | O70513 | 3.24 | 3.28E-04 | 3.86 | 1.97E-04 | 1.53 | 1.28E-02 |
| P08649 | Complement C4 (CO4) | P08649 | 2.24 | 3.62E-04 | 2.82 | 9.34E-04 | 3.15 | 1.02E-02 |
| P07151 | Beta-2-microglobulin (B2MG) | P07151 | 1.70 | 2.08E-03 | 2.05 | 1.10E-04 | 4.45 | 3.29E-03 |
| Q05820 | Putative lysozyme C-2 (LYZ2) | Q05820 | 1.65 | 3.71E-02 | 2.00 | 8.55E-04 | 2.29 | 1.09E-02 |
| Q6DGG1 | Protein ABHD14B (ABHD14B) | Q6DGG1 | 1.60 | 7.63E-03 | 1.53 | 5.75E-03 | 4.57 | 9.69E-03 |
| P16391 | RT1 class I histocompatibility antigen (HA12) | No | - | - | 3.93 | 6.55E-03 | 4.20 | 2.03E-03 |
| Q8JZQ0 | Macrophage colony-stimulating factor 1 (CSF1) | P09603 | - | - | 3.09 | 4.18E-03 | 3.00 | 2.88E-03 |
| Q1WIM1 | Cell adhesion molecule 4 (CADM4) | Q8NFZ8 | - | - | 2.20 | 1.27E-03 | 2.93 | 1.46E-02 |
| P02764 | Alpha-1-acid glycoprotein (A1AG) | P02764 | - | - | 1.92 | 1.01E-03 | 4.86 | 8.75E-03 |
| P80067 | Dipeptidyl peptidase 1 (DPP1) | P53634 | - | - | 0.60 | 6.48E-03 | 0.57 | 6.28E-04 |
| Q9JLJ3 | 4-trimethylaminobutyraldehyde dehydrogenase (TMABADH) | P49189 | - | - | 0.50 | 1.67E-02 | 0.19 | 2.70E-03 |
| Q711G3 | Isoamyl acetate-hydrolyzing esterase 1 homolog (IAH1) | Q711G3 | - | - | 0.33 | 1.17E-02 | 0.00 | 1.45E-03 |
| Q8R5M3 | Leucine-rich repeat-containing protein 15 (LRRC15) | Q8R5M3 | 6.20 | 9.19E-03 | 7.40 | 9.84E-04 | - | - |
| Q62638 | Golgi apparatus protein 1 (ESL1) | Q62638 | 4.75 | 6.67E-03 | 5.13 | 2.86E-03 | - | - |
| P02761 | Major urinary protein (MUP) | No | 0.39 | 3.89E-03 | 0.59 | 1.51E-02 | - | - |
| P62898 | Cytochrome c, somatic (CYCS) | P99999 | 3.33 | 1.63E-03 | - | - | 3.83 | 1.85E-03 |
| P01946 | Hemoglobin subunit alpha-1/2 (HBA1) | P69905 | 8.60 | 4.30E-02 | - | - | - | - |
| P20786 | Platelet-derived growth factor receptor alpha (PDGFRA) | P16234 | 3.38 | 8.97E-03 | - | - | - | - |
| P16310 | Growth hormone receptor (GHR) | P16310 | 2.71 | 4.42E-03 | - | - | - | - |
| P20762 | Ig gamma-2C chain C region | No | 2.25 | 2.45E-03 | - | - | - | - |
| P08753 | Guanine nucleotide-binding protein G(k) subunit alpha (GNAI3) | P08753 | 2.09 | 9.02E-03 | - | - | - | - |
| P08289 | Alkaline phosphatase (AP-TNAP) | P05186 | 1.57 | 9.02E-03 | - | - | - | - |
| P80204 | TGF-beta receptor type-1 (TGFR-1) | P80204 | 1.54 | 4.23E-04 | - | - | - | - |
| O88766 | Neutrophil collagenase | P22894 | 0.54 | 1.05E-02 | - | - | - | - |
| P84855 | Major urinary protein (MUP) | No | 0.44 | 8.37E-03 | - | - | - | - |
| P04218 | OX-2 membrane glycoprotein (MOX2) | P04218 | - | - | 3.25 | 1.06E-03 | - | - |
| P49134 | Integrin beta-1 (ITGB1) | P49134 | - | - | 3.14 | 5.33E-04 | - | - |
| Q9Z0J6 | Growth/differentiation factor 15 (GDF-15) | Q9Z0J6 | - | - | 3.13 | 2.51E-03 | - | - |
| P34058 | Heat shock protein HSP 90-beta (HSP 84) (HSP84) | P34058 | - | - | 3.11 | 2.50E-02 | - | - |
| Q9EPF2 | Cell surface glycoprotein MUC18 (MUC18) | Q9EPF2 | - | - | 3.08 | 6.82E-03 | - | - |
| Q9QWJ9 | Neuropilin-1 (NRP1) | Q9QWJ9 | - | - | 2.57 | 3.61E-03 | - | - |
| P07943 | Aldose reductase (AR) | P07943 | - | - | 2.44 | 1.13E-02 | - | - |
| P97603 | Neogenin (Fragment) (NEO1) | P97603 | - | - | 2.10 | 3.63E-04 | - | - |
| Q80WF4 | Transmembrane protein 132A (T132A) | Q80WF4 | - | - | 1.90 | 1.75E-02 | - | - |
| Q91XT9 | Neutral ceramidase (ASAH2) | Q91XT9 | - | - | 1.80 | 1.34E-02 | - | - |
| Q6MG71 | Choline transporter-like protein 4 (CTL4) | Q91XT9 | - | - | 1.71 | 7.43E-03 | - | - |
| Q63691 | Monocyte differentiation antigen CD14 (CD14) | Q63691 | - | - | 1.70 | 4.12E-03 | - | - |
| B5DFC9 | Nidogen-2 (NID-2) | B5DFC9 | - | - | 1.59 | 6.65E-03 | - | - |
| O35217 | Multiple inositol polyphosphate phosphatase 1 (MINPP1) | Q9UNW1 | - | - | 1.51 | 1.23E-03 | - | - |
| O88767 | Protein/nucleic acid deglycase DJ-1 (PARK7) | O88767 | - | - | 0.33 | 7.68E-03 | - | - |

**Figure 3.**
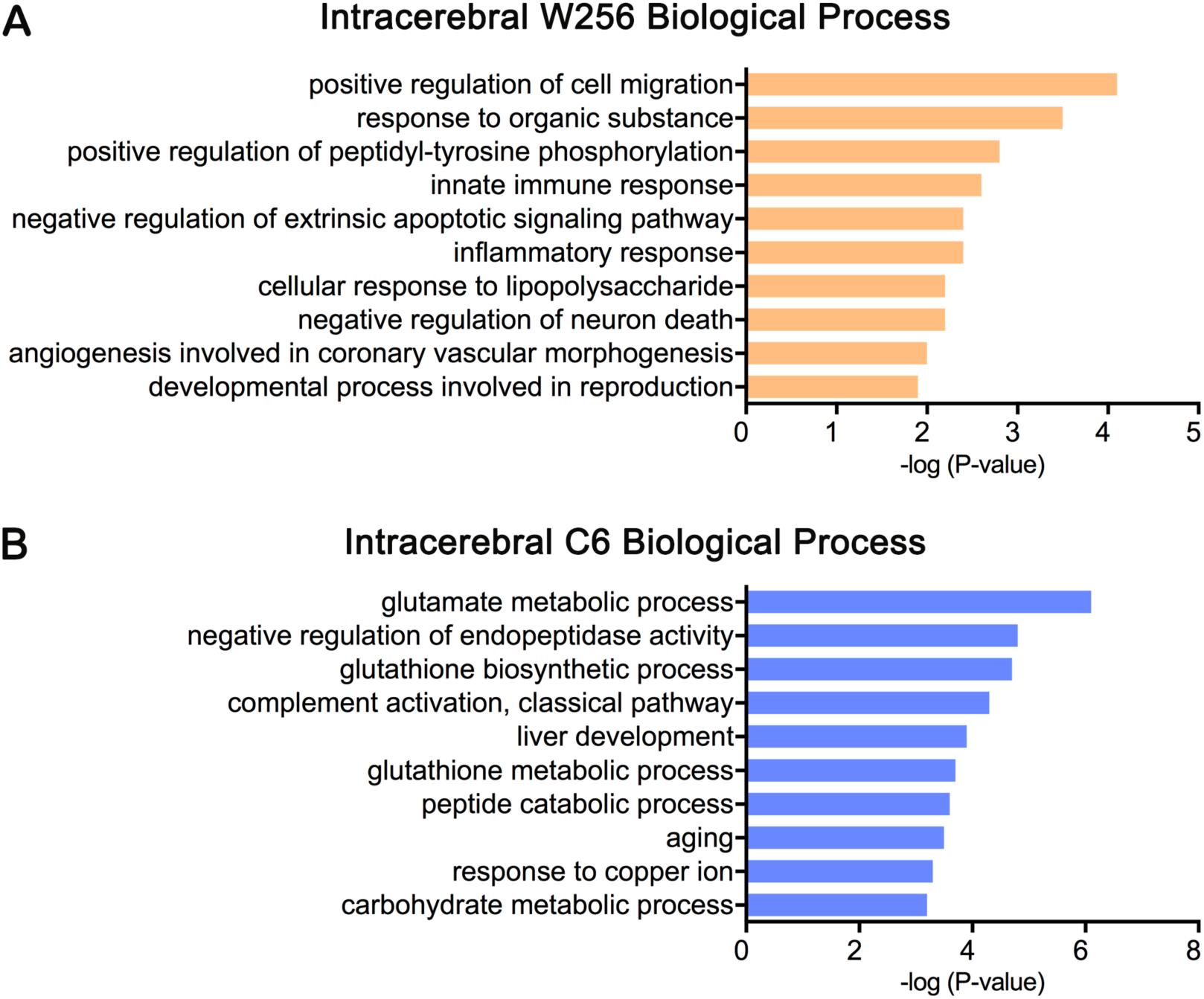
The 10 most significantly enriched biological processes. A) Early-stage changed proteins in the intracerebral W256 model; B) Early-stage changed proteins in the intracerebral C6 model.

Interestingly, tumors did not grow in four rats after injection of highly malignant W256 cells. The urine samples at three time points (days 5, 8 and 10) from these four tumor-rejecting rats were also analyzed to investigate whether urinary protein levels can reflect some currently unclear changes in the body after W256 injection. The same analysis methods as above were used, and the results showed that several proteins were altered on days 5, 8 and 10, suggesting that the urinary proteome was changed in the tumor-rejecting rats. We hypothesized that the urinary protein changes may be related to the eradication of a few inoculated tumor cells by the body’s immune system, in addition to the recovery from surgery. For example, some proteins, such as TIMP1, GRP78, NUCB2 and CALB1, were found to be significantly changed only in the tumor-rejecting rats (Figure 4). TIMP1 is an inhibitor of matrix metalloproteinases, a matrix-degrading enzyme that facilitates tumor cell dissemination[12]. High serum levels of TIMP-1 in patients with a variety of cancers have been associated with poor prognosis in a variety of cancer patients.

**Figure 4.**
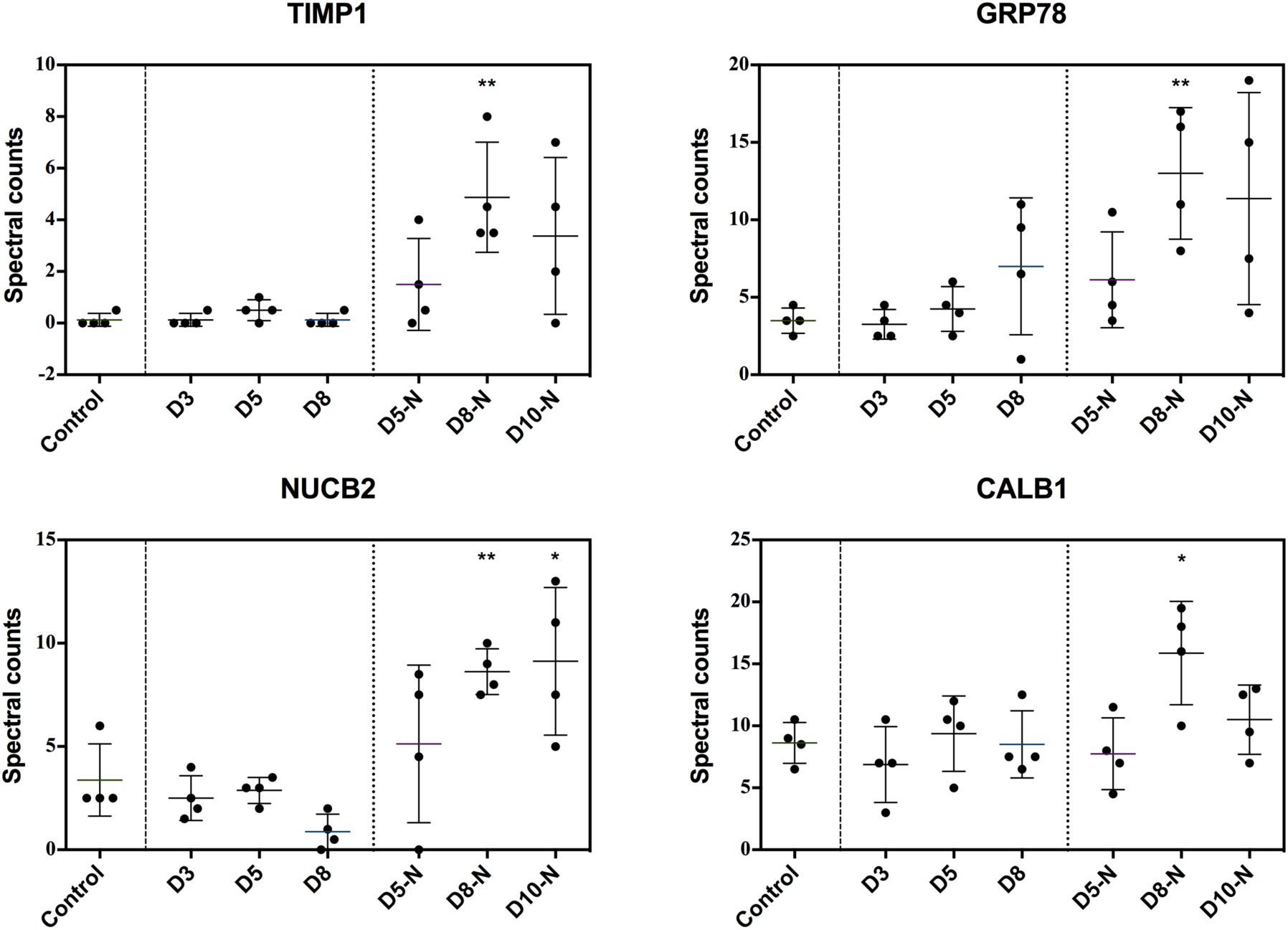
Relative quantitation of four changed urinary proteins that significantly changed only in tumor-rejecting rats (D5-N, D8-N and D10-N), P <0.05 (*), P<0.01 (**).

Additionally, urinary TIMP-1 was found to be a predictor of few cancers, such as renal carcinoma[13], bladder cancer[14] and pancreatic malignancies[15]. GRP78 expressed on the surface of tumor cells is associated with proliferation and metastasis. The presence of cancer-related proteins that were significantly changed in tumor-rejecting rats but not in tumor rats suggests that tumor cells were indeed injected into the brain and eventually eliminated due to the high immunity.

### Urinary proteomic comparative analysis of different tumor models

As shown in Figure 5, comparison of all differentially expressed proteins indicated that the proportion of overlapping proteins was small, and more than half of the differential proteins in each of the three models were unique. The details of the differential proteins in the three models are shown in Table S2.

**Figure 5.**
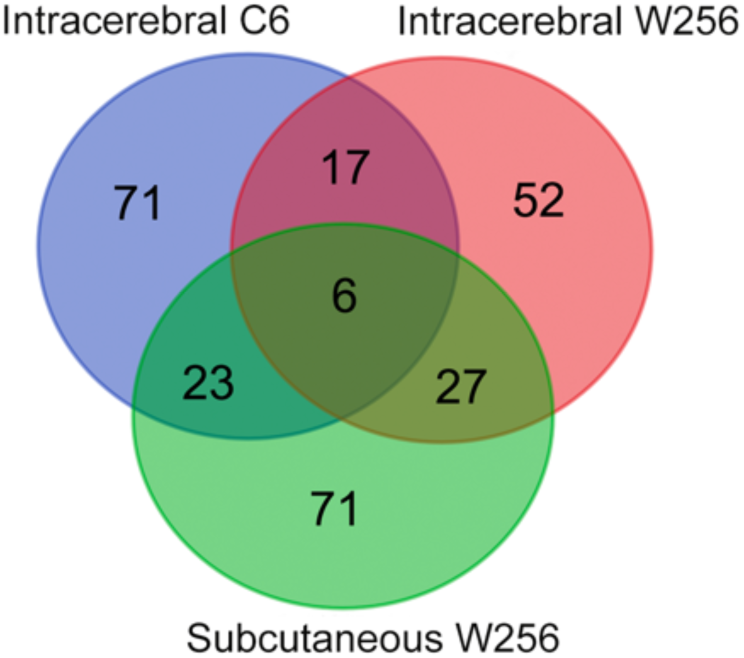
Venn diagram indicating the overlapping differential proteins in urine samples of the subcutaneous W256, intracerebral W256 and intracerebral C6 models.

Comparing the differential proteins of the intracerebral C6 and intracerebral W256 models showed only 23 overlapping proteins. Ten of the 23 proteins were identified in the early stages of both models, and among them, the GDF15, CTL4 and PARK7 proteins were associated with nervous system function. In contrast to the intracerebral W256 model, in which a large number of differential proteins were identified at the late time point, most differential proteins could be identified at the early stages in the intracerebral C6 model. The 101 early-stage differential proteins in the intracerebral C6 model were also annotated in the GO biological process analysis (Figure 3B). The main significantly enriched biological processes were different from those of the intracerebral W256 model. For example, glutamate metabolic process was significantly overrepresented in the intracerebral C6 group. Glutamate is the major excitatory neurotransmitter in the central nervous system, and abnormally increased glutamate levels destroy normal tissues. Glioma cells produce glutamate, which promotes tumor invasion through destruction of the surrounding brain tissue[16]. These results suggest that this effect may be due to the differences in invasion abilities and patterns of W256 and C6 cells. W256 cells in brain tissue showed expansive growth with distinct borders between tumors and the surrounding brain parenchyma, while C6 cells diffusely infiltrated the brain parenchyma and displayed strongly invasive extension[8].

After comparing the differences in urinary proteomes between the intracerebral W256 model and the subcutaneous W256 model, we found that only less than a third of the proteins overlapped, and the difference was greater in the late stage of the tumor. However, in the early stages, more tumor metastasis-associated proteins were identified in both the intracerebral and subcutaneous W256 models, such as B2MG, A1AG, LG3BP and CSF1, but showed no significant changes in the intracerebral C6 model. This finding suggests that a greater number of similar urine changes will occur in the early stages when the same tumor cells are grown in different organs.

### Urinary candidate biomarker validation and analysis

To find more reliable urinary differential proteins associated with intracerebral tumors, the same 4 urine samples in control rats that were previously used and 12 urine samples from three time points (days 3, 5 and 8) in another four tumor rats were chosen for validation. There were 20, 17 and 66 changed proteins on days 3, 5, and 8, respectively (fold change ≥1.5 or ≤0.67, P < 0.05, Table S3). The commonly identified proteins that significantly changed in the eight tumor rats are listed in Table 2.

**Table 2.**
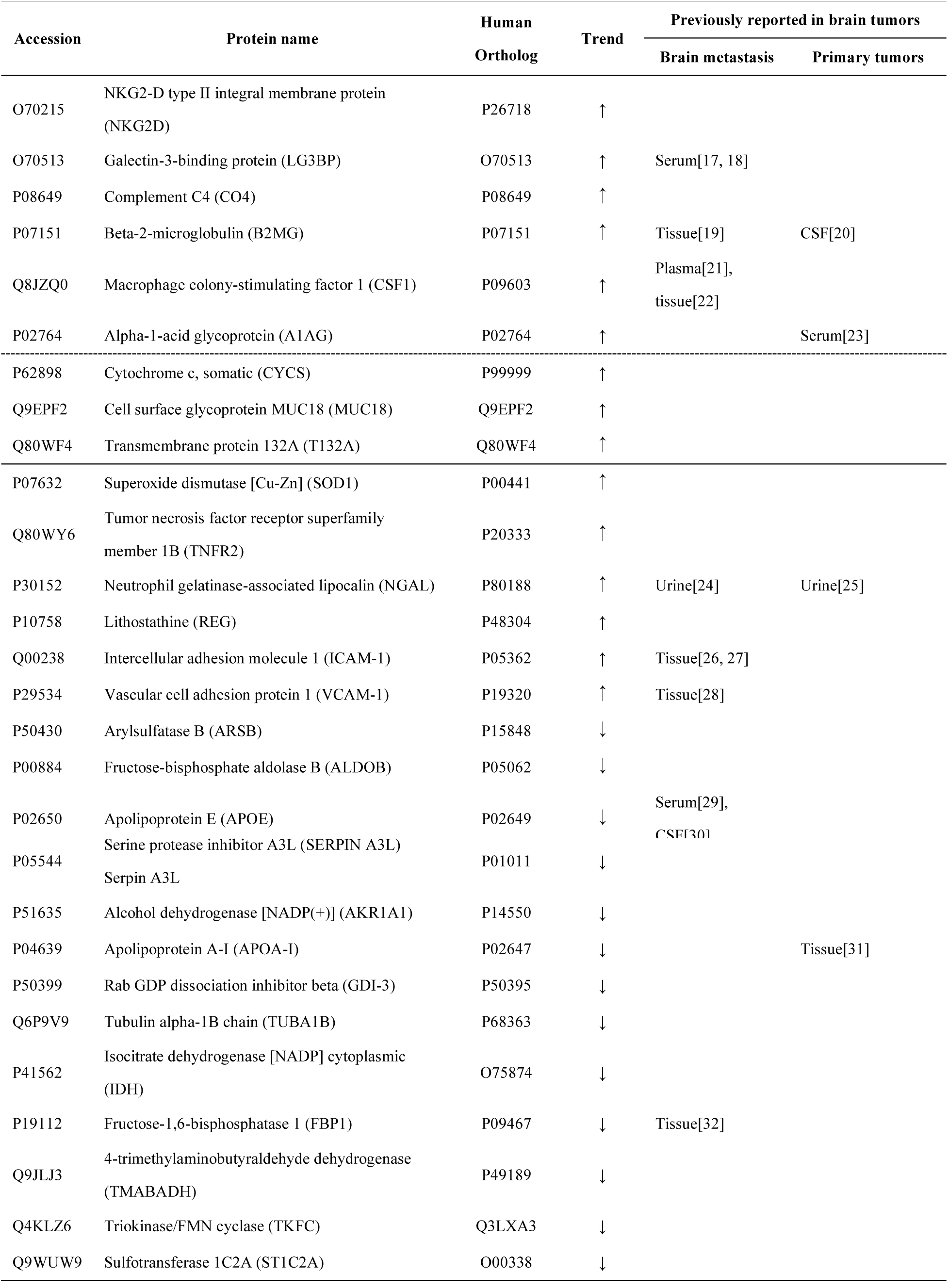
**Differential proteins identified in eight intracerebral W256 rats.**

Among 43 early-stage changed proteins identified above, nine proteins (NKG2D, LG3BP, CO4, B2MG, CSF1, A1AG, CYCS, MUC18, T132A) that had human orthologs were also identified in the early stages of the other four rats, and the first 6 proteins continued to change differentially on day 8. Among them, CO4, B2MG and A1AG enable early detection of cancer according to Wu *et al*.[7]. B2MG has been reported to be a potential marker of meningioma[20] and brain metastases in breast cancer[19]. A1AG is a serum acute phase response protein with immunomodulatory effects and can also be used as a sensitive indicator of inflammation levels and tissue damage levels[33]. Additionally, LG3BP is overexpressed in the serum or tissue of various cancer patients and plays roles in cancer cell aggregation and metastasis[17, 18]. Overexpressed CSF1 with high plasma levels is associated with poor prognosis of malignant tumors, such as breast cancer[21], and CSF1 has also been reported to be expressed in brain metastases of lung cancer and breast cancer[22]. Furthermore, NKG2D acts as an activating receptor for killer cells (NK), which can be induced by hyperproliferation and transformation and by pathogen infection, providing therapeutic targets for the treatment of infectious diseases, cancer and autoimmune diseases[34].

On day 8, the tumor rats lost weight and tumors were detectable on MRI. Among the 75 changed proteins mentioned above, 25 that had human orthologs were also identified in the other four rats on day 8. In addition to NKG2D, LG3BP, B2M, CSF1 and A1AG, several proteins had been reported to be differentially expressed in serum, urine or brain tissue of brain metastasis or primary tumors. For instance, NGAL is a secreted glycoprotein that can be used as a biomarker for cancer diagnosis and prognosis[35], and increased levels have been reported in the urine of patients with cancer, such as brain tumors[25] and breast cancer[24]. ICAM-1, a cell surface glycoprotein that is mainly expressed in nervous tissues and tumors, has been reported to be upregulated in brain tissue of patients with lung cancer or breast cancer[26] and two breast cancer brain metastasis mouse models[36]. APOE is mainly involved in the regulation of lipid metabolism. Recent studies have shown that APOE can play a role in inhibiting cancer metastasis and act as a serum biomarker for the assessment of lung cancer metastasis[29]. Additionally, APOE was decreased in the cerebrospinal fluid of breast cancer meningeal metastasis patients[30]. VCAM-1 in brain metastasis-related cerebral vessels was upregulated significantly with the progression of tumor, and this change was validated in both brain-transferred mice and human brain tissue, indicating the important roles of VCAM-1 in the establishment and progression of brain metastasis[28].

Urine, a new source of sensitive biomarkers for noninvasive analysis, has been shown to provide clues for the early diagnosis of disease in a variety of animal models in our previous work[37-42]. The W256 breast carcinoma cell line has been used for a rat model of brain metastases that is similar in histopathology to patients with brain metastases[43-46]. Direct intracerebral injection of tumor cells can lead to tumor cell growth in the brain, which is the most well-controlled and reproducible method for establishing animal models of brain metastases[47, 48]. Therefore, these proteins or combinations that have changed in urine without clinical manifestations and MRI changes may provide clues for the early diagnosis of brain metastases as the intracerebral W256 model mimics the local tumor growth process of brain metastases. Additionally, the differential diagnosis between gliomas and single brain metastases with an unclear primary disease history is clinically difficult because of the similar clinical and MRI manifestations, and there are differences in diagnosis and treatment strategies among these two diseases. Urinary proteomic comparative analysis among the intracerebral W256, subcutaneous W256 and intracerebral C6 models revealed that urine proteomics may provide some new clues for definitive diagnosis.

## Conclusion

In this study, several urinary differential proteins were identified on days 3 and 5 before the appearance of obvious abnormalities on MRI in an intracerebral W256 model. Nine proteins, including LG3BP, CSF1 and NKG2D, known to play important roles in the proliferation and metastasis of cancer, were changed significantly. When tumors were detected by MRI, twenty-five differential proteins were identified in the urine of all 8 tumor rats, which may be useful for monitoring the progression of tumors and even the differential diagnosis of primary brain tumors and brain metastases. Additionally, the urine proteome may provide a new direction for the diagnosis of primary tumor metastases and metastatic organs. Given the small number of model animals in this study, a larger number of clinical samples are needed for verification in future research.

## Supporting information

## Acknowledgements

This research was supported by the National Key Research and Development Program of China (2018YFC0910202, 2016YFC1306300), Beijing Natural Science Foundation (7172076), Beijing cooperative construction project (110651103), Beijing Normal University (11100704) and Peking Union Medical College Hospital (2016-2.27).

